# Hypometabolism in Autism Spectrum Disorder: Insights from Brain and Blood Transcriptomics

**DOI:** 10.1101/2024.11.18.624092

**Authors:** Rami Balasubramanian, Debayan Saha, Ananya Arun, PK Vinod

## Abstract

Autism Spectrum Disorder (ASD) is a neurodevelopmental condition characterized by challenges in social communication, repetitive behaviors, and restricted interests. Recent research has emphasized the importance of metabolic dysfunctions in the pathophysiology of ASD. This study investigates metabolic alterations associated with ASD by analyzing transcriptomic data obtained from the prefrontal cortex (bulk tissue and single-nucleus) and data from peripheral blood mononuclear cells (PBMC). We assessed the metabolic activity of each patient based on gene expression profiles, revealing significant downregulation of vital metabolic pathways, including glycolysis, the tricarboxylic acid (TCA) cycle, and oxidative phosphorylation, indicative of hypometabolism. Our analysis also highlighted dysregulation in lipid, vitamin, amino acid, and heme metabolism, which may contribute to the neurodevelopmental delays associated with ASD. Cell-specific metabolic activities in the ASD brain showed altered pathways in astrocytes, oligodendrocytes, excitatory neurons, and interneurons. Furthermore, we identified critical metabolic pathways and genes from PBMC gene expression data that distinguish ASD patients from typically developing individuals. Our findings demonstrate a consistent pattern of metabolic dysfunction across brain and blood samples. This research provides a comprehensive understanding of metabolic alterations in ASD, paving the way for exploring potential therapeutic strategies targeting metabolic dysregulation.

## Introduction

Autism spectrum disorder (ASD) is a neurodevelopmental condition characterized by core deficits in social communication and interaction along with restricted and repetitive behaviors. According to the CDC, it affects approximately 1 in 36 children and presents a wide range of severity and symptoms [1]. Despite common symptoms, ASD displays significant variability, both in severity and genetic factors, making it challenging to identify specific causes and develop targeted treatments. Recent research suggests a link between ASD and specific metabolic dysfunctions, indicating that aberrations in metabolic pathways may contribute to the neurobiological underpinnings of the disorder. Mitochondrial dysfunction has been observed in some individuals with ASD [2]. This dysfunction may disrupt critical neuronal processes and contribute to the development of ASD symptoms. Studies have shown potential deficiencies in folate metabolism or elevated levels of folate receptor antibodies in some individuals with ASD [3]. Conditions like phenylketonuria, a disorder of amino acid metabolism, and creatine deficiency syndromes have been identified in some individuals with ASD, suggesting broader metabolic imbalances [4]. In an earlier study, we mapped the transcriptomic changes associated with neuropsychiatric conditions: bipolar disorder (BD), schizophrenia (SCZ), and depression (MDD), providing insights into the metabolic alterations in these conditions [5].

Given the complex interplay between metabolism and neurodevelopment, there is a growing interest in employing omics data to investigate metabolic dysregulation in ASD, both at the bulk tissue and single-cell levels. Gene expression patterns of metabolic genes can be inferred using transcriptomics data of patients, which can help to elucidate the metabolic landscape of ASD. Metabolic changes can be analyzed at different levels, including pathway, metabolite, and flux levels, each offering unique insights into cellular metabolism (Hrovatin et al., 2022). Pathway-level analysis focuses on studying the activity of specific metabolic pathways by evaluating gene expression data related to enzymes and reactions within those pathways. This method is valuable for understanding how pathways are activated or suppressed in response to various conditions, helping to identify critical pathways involved in disease conditions. The metabolite-level analysis involves identifying metabolites associated with differentially expressed genes and mapping to the associated pathways to understand how gene expression alterations influence the overall metabolic network [7]. Flux-level analysis uses computational models like Flux Balance Analysis (FBA) to predict the flow of metabolites through metabolic pathways, providing a quantitative and dynamic view of metabolism [8]. These methods can also be applied to single-cell transcriptome data to uncover cellular heterogeneity and to understand the metabolic profiles of distinct cell populations in the context of ASD [6].

In our study, we explored the metabolic landscape of ASD patients by analyzing transcriptomic data from brain and blood samples (Figure 1). We assessed the metabolic pathway activity for each sample by scoring the individual pathways of the human metabolic network. This approach allowed us to investigate differences in pathway activity between ASD and control samples. We also identified metabolites that are associated with significant transcriptional changes. To address cellular heterogeneity, we analyzed the single-nucleus RNA-Seq data from the ASD brain, allowing us to capture metabolic variation at the cellular level. By analyzing bulk tissue and single-nucleus transcriptomes of key brain regions and peripheral blood, we shed light on the hypometabolic states of ASD. We also identified potential metabolic genes from peripheral blood transcriptome profiles that may act as diagnostic markers to distinguish ASD from typical development. Our comprehensive analysis sheds light on the metabolic changes in both the brain and peripheral blood of individuals with ASD and holds the potential for improving diagnostic and therapeutic strategies.

**Figure 1.**
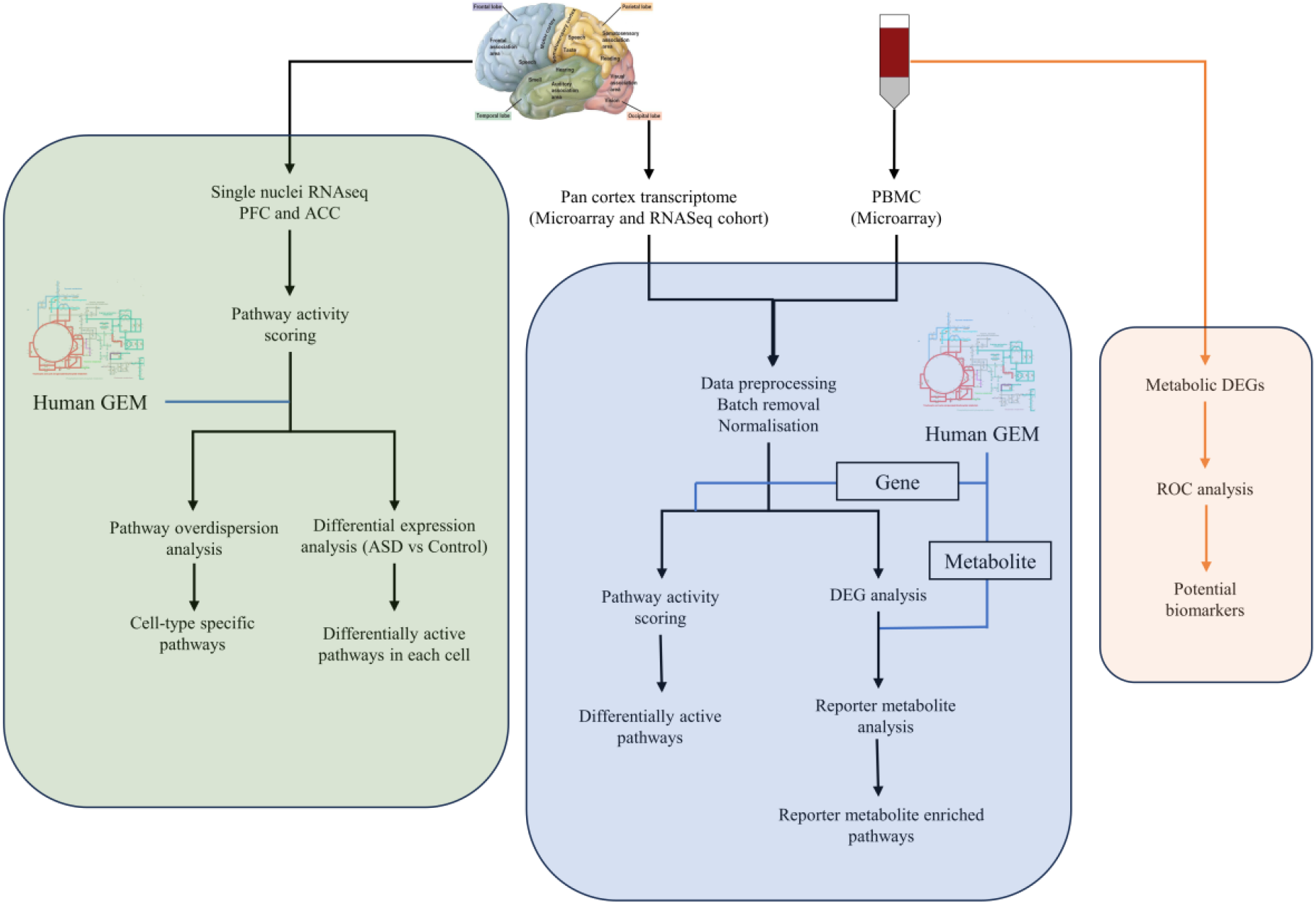
Overall workflow to investigate the differential metabolic activity of ASD using brain tissue and peripheral blood transcriptomes.

## Methods

### Dataset description and processing

We analyzed transcriptome data from post-mortem brain samples and peripheral blood mononuclear cells (PBMCs) to understand metabolic activity changes associated with ASD. Bulk-tissue and single-nucleus brain transcriptome data, along with PBMC microarray data, were obtained from multiple studies to perform a comprehensive analysis (Table S1). Pre-processed pan-cortex bulk-tissue transcriptomic data from Gandal et al. (2018) was used, which includes microarray data from Brodmann areas (BA) BA9, BA41 (GSE28521), BA9/46 (GSE28475), and BA41/42 (Garbett et al. 2008), as well as pan-cortical RNA-seq data from BA4/6, BA38, BA7 and BA17 (syn11242290) (Table S1). The ASD cortex microarray datasets were normalized based on the array platform and processed independently. Metadata for the microarray cohort was generated by integrating the datasets and correcting batch effects using COMBAT in R [9]. Differentially expressed genes between ASD and control were identified using a linear mixed-effects model with the *nlme* package in R [10]. Normalized and log-transformed pan-cortical RNA-seq data were also used to compare and validate the results obtained from the microarray data. Differential gene expression analysis of the RNA-seq data was performed using the *limma* R package [11]. For the blood-based analysis, normalized and batch-corrected microarray data of PBMCs (GSE18123) from ASD and typically developing children were obtained from GEMMA (https://gemma.msl.ubc.ca/home.html) [12]. Differential gene expression analysis of PBMC data was also performed using the *limma* R package[11].

We obtained single-nucleus RNA-seq (snRNA-seq) data from 48 post-mortem tissue samples of the prefrontal cortex (PFC) from 13 patients with ASD and 10 control subjects (https://cells.ucsc.edu/?ds=autism) [13]. The following criteria were applied to filter cells and genes for the downstream analysis: cells expressing ≥500 genes, with less than 5% mitochondrial and ribosomal transcript fractions, and genes expressed in more than five cells [13]. Matrices were normalized to total UMIs per nucleus and log-transformed. For each cell cluster, we identified differentially expressed pathways [11].

### Pathway activity

Pathway activity score for each sample was calculated using singscore, a rank-based metric that measures gene set enrichment of metabolic pathways from the Human-GEM, a generic human genome-scale metabolic network [14]. The gene signatures of metabolic pathways were undirected, and the absolute median-centered rank was computed based on gene expression (equation 1).

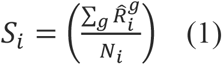

*S*_*i*_ is the score for sample *i* in relation to the pathway gene set, and 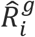 is the absolute, median-centered rank of gene *g* in the gene set. *N*_*i*_ is the number of genes in the gene set. 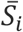 is the normalized score for sample *i* in relation to the pathway gene set (equation 2).

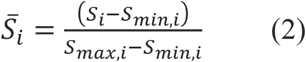

*S*_*max,i*_ and *S*_*min,i*_ are the theoretical minimum and maximum mean ranks obtained from the arithmetic series expansion (equation 3 and 4).

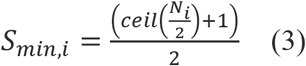

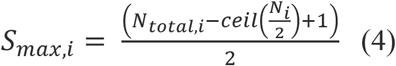

The pathway activity is scored for individual samples, and the resulting scores range between 0 and 1. The differential metabolic activity between ASD and control was analyzed using the *limma* R package. The hierarchical clustering of the samples was performed with the differentially altered pathways (adjusted p-value < 0.05).

For single-nucleus RNA-seq data, metabolic activity was analyzed using the PAGODA2 R package, which scores cells based on the first weighted principal component. Pathway overdispersion analysis was performed with default parameters to identify cell heterogeneity [15]. The scores of the significantly over-dispersed pathways were then used for t-distributed Stochastic Neighbor Embedding (t-SNE), enabling visualization of the heterogeneous metabolic activity across cells.

### Reporter metabolite analysis

Reporter metabolites analysis identifies metabolites in the neighborhood of enzymes of significant transcriptional changes by integrating differential gene expression into the metabolic network. We used the Human-GEM model as the metabolic network [16], and mapped the differentially expressed gene (log fold change of genes and the p-value from the differential expression analysis) to the metabolic network using Raven 3.0 [7]. The p-value of the genes were converted into Z-scores (*Z*_*i*_) using the inverse normal cumulative distribution function (CDF). Metabolites were assigned a Z-score (*Z*_*metabolite*_) by aggregating the Z scores of the ‘k’ neighboring genes (equation 5).

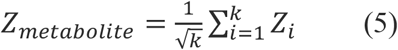

The mean (*μ*_*k*_) and standard deviation (*σ*_*k*_) of the aggregated Z scores, generated by sampling 10,000 sets of k enzymes from the network, were used to adjust the *Z*_*metabolite*_ scores for the background distribution (equation 6).

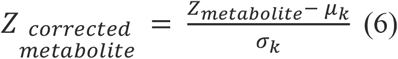

The corrected Z-scores were converted to p-values using CDF, and the metabolites with p-values less than 0.05 were considered reporter metabolites. We identified the metabolites that are associated with up and downregulated genes. To find the pathways that are enriched with reporter metabolites, we performed the hypergeometric enrichment test with the pathways and corresponding metabolites from the Human-GEM network.

### ROC analysis

To identify the most effective diagnostic markers for ASD, we analyzed differential metabolic pathways and genes using receiver operating characteristic (ROC) analysis. The pathway scores from ASD-PBMC samples, derived using singscore, were used for ROC analysis. The pROC R package was used to calculate the area under the ROC curve (AUC), enabling the identification of the top features (pathways and genes) with high AUC values. The top features were validated using PBMC and leukocyte microarray transcriptome data (Table S1).

### Cross-disorder analysis

We also investigated metabolic activity across ASD conditions, including autism, Asperger’s disorder (AD), and Pervasive Developmental Disorder-Not Otherwise Specified (PDD-NOS), using blood-based transcriptome data. Metabolic pathway activity scores for each sample were calculated using singscore, and pathways showing differential expression compared to controls were identified across ASD conditions. The adjusted p-values from the differential expression analysis were transformed to -log10 of adjusted p-values. These transformed values served as the basis for the PCA biplot analysis to identify and visualize metabolic differences among the conditions. Additionally, we also performed a comparative study with other major neuropsychiatric disorders, such as schizophrenia (SCZ), bipolar disorder (BD), and major depressive disorder (MDD), using brain cortex microarray data (Table S1) [17].

## Results

### Differential metabolic activity in the ASD brain

To investigate metabolic variations in ASD samples compared to controls, the expression of metabolic genes from the Human-GEM network was analyzed. Differential expression analysis was performed on both the ASD microarray and RNA-Seq cohorts. Principal component analysis using differentially expressed genes (adjusted p-value < 0.01) revealed considerable variability among samples in both cohorts. Figure 2 illustrates the separation of ASD samples from controls. However, some ASD samples overlap with controls, highlighting the heterogeneity within ASD cases. This variability underscores the need for a more detailed sample-level analysis of metabolic activity.

**Figure 2.**
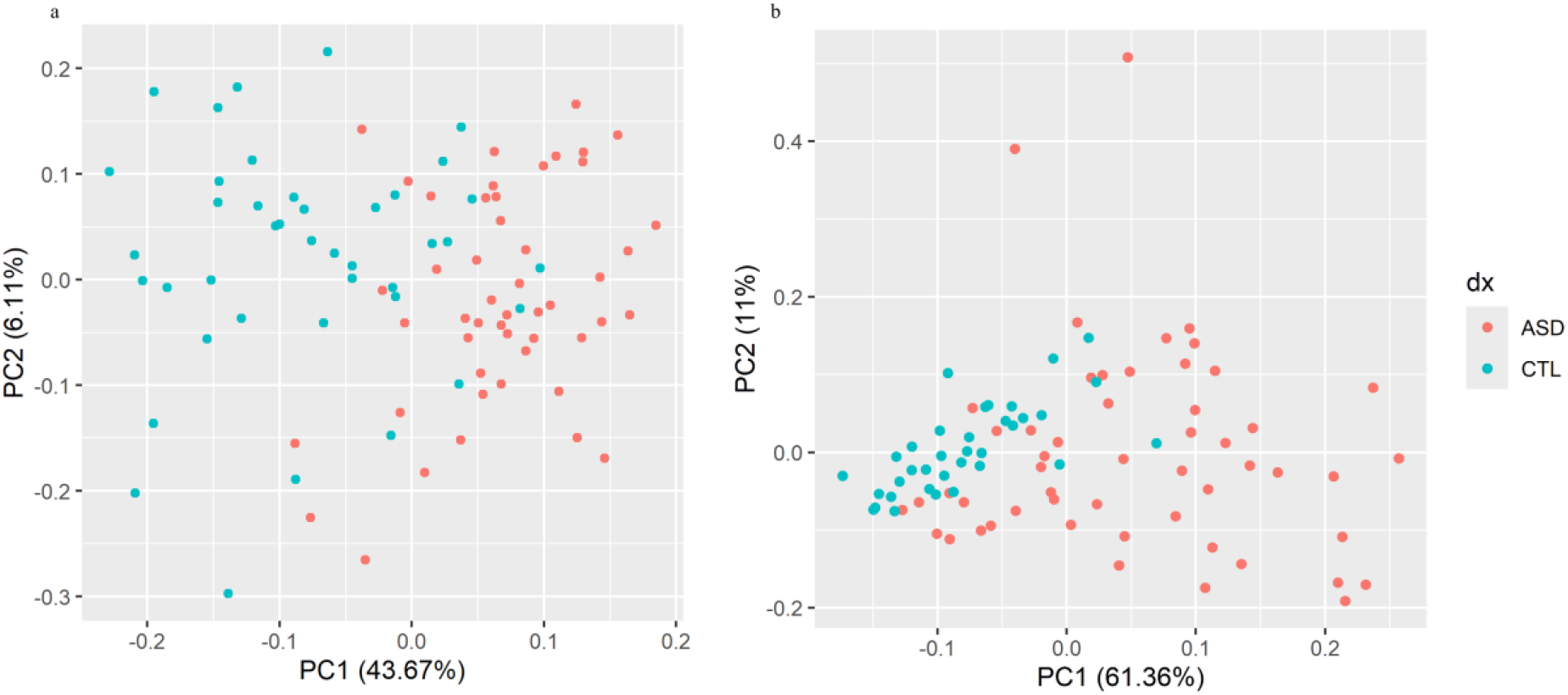
Principal components of differentially expressed metabolic genes showing variation between ASD and Control in the microarray cohort (a) and RNA-seq cohort (b).

We aggregated the genes based on metabolic pathways and calculated the pathway activity by scoring each patient using singscore, a gene-set scoring method (see methods). These scores were used to identify significantly altered metabolic pathways between ASD and control groups (adjusted p-value < 0.05) (Table S2, S3). Hierarchical clustering of samples based on the scores of significant metabolic pathways revealed variations in metabolism both within ASD and between the ASD and control groups (Figure 3).

**Figure 3.**
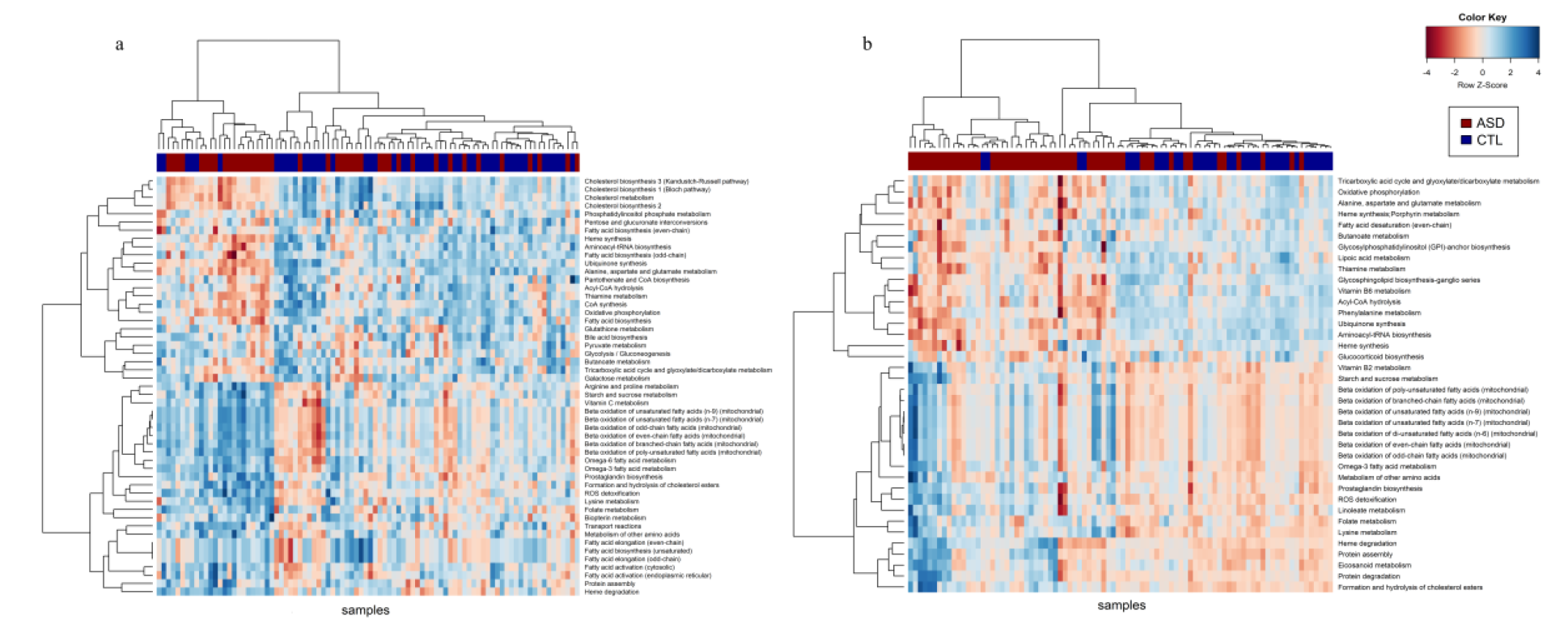
Hierarchical clustering of samples based on pathway activity score of differentially expressed metabolic pathways in the microarray cohort (a) and RNA-Seq cohort (b).

In both the microarray and RNA-Seq cohorts, we observed consistent downregulation of pathways related to glycolysis, the TCA cycle, and oxidative phosphorylation in ASD. Additionally, lipid metabolism, amino acid metabolism, glycan metabolism, and vitamin and cofactor metabolism pathways showed significant dysregulation. Cholesterol biosynthesis and metabolism pathways were markedly downregulated, while omega-6 and omega-3 fatty acid metabolism, prostaglandin biosynthesis, and linoleate metabolism pathways were upregulated. Pathways involved in glycan biosynthesis, such as glycosphingolipid biosynthesis (ganglio series) and Glycosylphosphatidylinositol (GPI)-anchor biosynthesis, also exhibited reduced activity in ASD. Alanine, aspartate, glutamate metabolism, and phenylalanine metabolism were downregulated, whereas arginine, proline, and lysine metabolism were upregulated in ASD. Folate metabolism was consistently elevated, while thiamine and vitamin B6 metabolism showed reduced activity. Furthermore, heme synthesis and porphyrin metabolism were downregulated, while heme degradation pathway was upregulated in ASD.

The differentially expressed metabolic genes were mapped onto the metabolite-enzyme bipartite network (Human-GEM) to identify metabolites associated with significant transcriptional changes across both cohorts (p-value < 0.05) (Table S4, S5). Metabolic pathways enriched with the altered reporter metabolites were determined using a hypergeometric enrichment test. Metabolites linked to glycolysis, the TCA cycle, oxidative phosphorylation, and lipid metabolism, including cholesterol metabolism, showed reduced activity (Figure 4). Significant alterations were observed in metabolites associated with alanine, aspartate, and glutamate metabolism, aromatic and branched-chain amino acid metabolism, arginine and proline metabolism, heme metabolism, and glutathione metabolism (Figure S1). The changes observed in reporter metabolite analysis align with the pathway activity in the individual samples.

**Figure 4.**
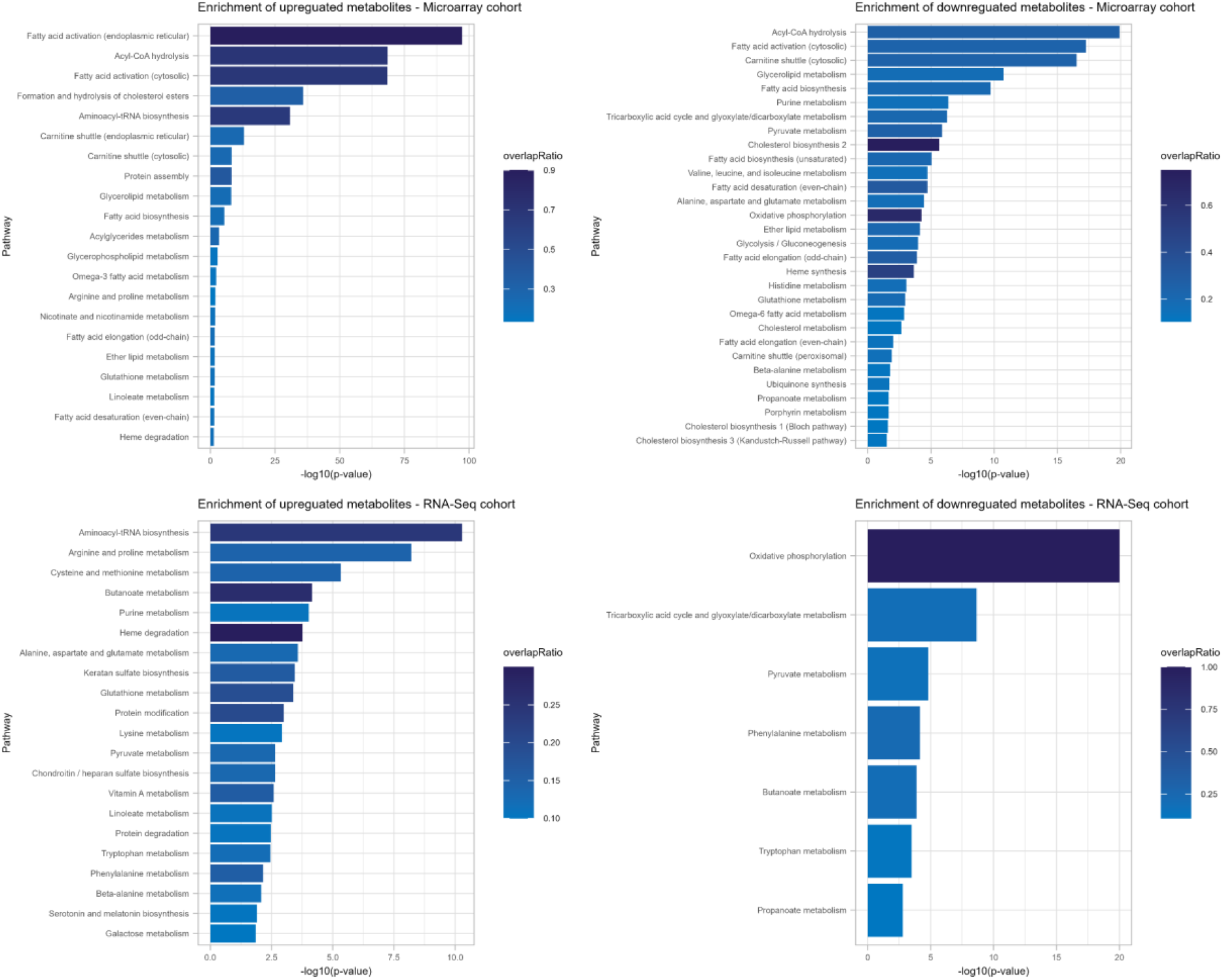
Pathways significantly enriched with reporter metabolites that are altered in both ASD brain cohorts.

### Distinct metabolic alterations in ASD compared to other neuropsychiatric disorders

To compare ASD with other neuropsychiatric disorders (SCZ, BD, and MDD), we performed a cross-disorder analysis by examining the differentially expressed pathways across various conditions in the brain cortex and identifying both overlapping and unique metabolic changes specific to ASD. This analysis revealed that ASD exhibits distinctive and extensive metabolic changes compared to other major psychiatric conditions (Table S6-S8). Although ASD shared certain metabolic pathways with SCZ and BD, it showed minimal overlap with MDD (Figure S2). Pathways such as oxidative phosphorylation, cholesterol metabolism, alanine, aspartate, and glutamate metabolism, sphingolipid, and galactose metabolism were downregulated in ASD, SCZ, and BD. In contrast, folate metabolism and lysine metabolism were upregulated. Interestingly, glucocorticoid biosynthesis showed contrasting patterns between ASD and MDD. In MDD, this pathway was upregulated, potentially indicating increased stress hormone synthesis linked to mood regulation and stress responses. In contrast, ASD showed downregulation of glucocorticoid biosynthesis, which might reflect differing neurobiological responses in ASD compared to MDD.

### Cell-specific metabolic activity

Investigating cell-specific metabolic activity is crucial for understanding the distinct metabolic functions of various cell types. In this study, we analyzed snRNA-seq data from the post-mortem brain cortex for cell-specific metabolic activities. We quantified pathway activity for each cell and identified 29 metabolic pathways that were significantly over-dispersed across the different cell types. t-SNE analysis of pathway activity scores revealed cellular heterogeneity, with distinct clusters of excitatory neurons, interneurons, maturing neurons, NRGN-expressing neurons, and various glial cells (Figure 5a, b). Astrocytes exhibited elevated expression of pathways related to protein assembly and degradation and amino acid metabolism, including alanine, aspartate, glutamate, and branched-chain amino acid (BCAA) metabolism, compared to other cell types (Figure 5c). Astrocytes are crucial to neurotransmitter balance, managing glutamate through the glutamate-glutamine cycle and aiding in branched-chain amino acid metabolism to support glutamate synthesis.

**Figure 5.**
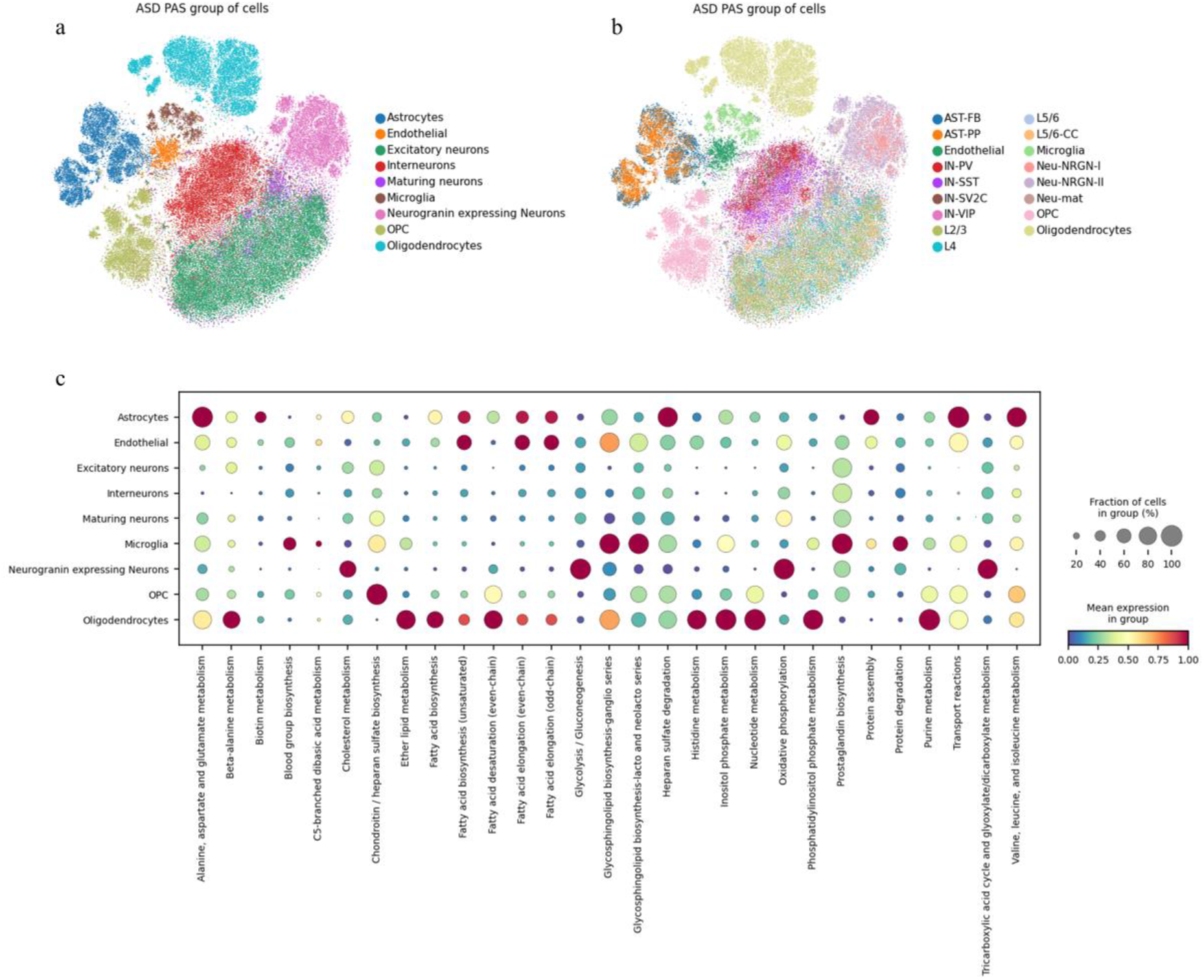
Cell-specific metabolic activity of ASD brain: t-SNE of cells defined by metabolic pathway activity score and colored based on particular cell group (a, b). Dot plot explaining the cell-specific expression of metabolic pathways (c).

Glycosphingolipid biosynthesis (ganglio-series, lacto and neolacto series) and prostaglandin biosynthesis were significantly upregulated in microglia. Glycosphingolipid biosynthesis in the brain is crucial for cell signaling, cell-to-cell communication, and immunomodulation. Gangliosides play an essential role in modulating microglial inflammatory responses. Fatty acid metabolic pathways are more specific to oligodendrocytes, which support neurons and signal transmission in the CNS by wrapping nerve fibers with a lipid-rich myelin membrane. Both fatty acid synthesis and fatty acid breakdown through β-oxidation occur in oligodendrocytes [18]. Energy-associated metabolic pathways, including glycolysis/gluconeogenesis, the TCA cycle, and oxidative phosphorylation, were more active in Neurogranin (NRGN) expressing neurons. NRGN is a protein that is critical in synaptic function and plasticity.

We observed that these 29 overdispersed pathways exhibited cellular heterogeneity without a significant ASD-specific effect, likely due to the masking of disease-specific signals by cell overdispersion. To address this, we performed pathway activity scoring and differential activity analysis for each cell type to identify pathways significantly altered in ASD compared to control. Our study showed consistent downregulation of oxidative phosphorylation across all cell types in ASD (Table S9-S13). In excitatory and interneurons, O-glycan metabolism, chondroitin/heparan sulfate biosynthesis, and heparan sulfate degradation were upregulated in ASD, while the TCA cycle was downregulated (Table S9, S10). Furthermore, aromatic amino acid and serotonin-melatonin biosynthesis were upregulated in excitatory neurons (Figure 6). These findings indicate that excitatory and interneurons show more disease-level changes in ASD conditions, consistent with findings reported in [19]. In astrocytes, glycolysis/gluconeogenesis and cholesterol metabolism were significantly downregulated. Cholesterol metabolism was also downregulated in oligodendrocyte progenitor cells (Table S11-S13).

**Figure 6.**
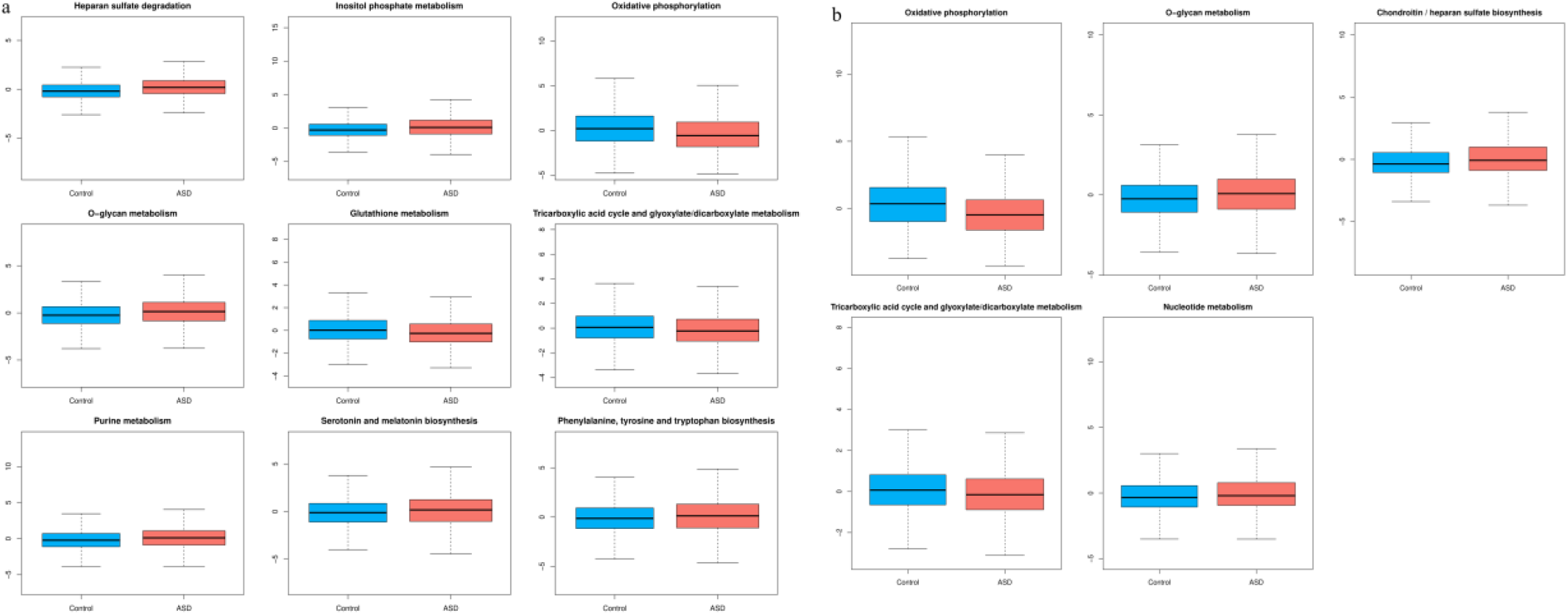
Differentially altered pathways in excitatory neurons (a) and interneurons (b).

### Differential metabolic pathway activity in peripheral blood samples of ASD

Building on our brain transcriptome analysis that uncovered ASD-related metabolic changes, we aimed to investigate whether similar alterations are present in the blood samples. We analyzed the PBMC transcriptome data of typically developing controls and ASD individuals. Metabolic pathway activity scores were calculated, and significantly altered pathways in ASD were identified (Figure S3). Our results indicate that several key metabolic pathways, including aromatic amino acid metabolism, arginine and proline metabolism, alanine, aspartate, and glutamate metabolism, and glutathione metabolism, were downregulated in ASD. Chondroitin/heparan sulfate biosynthesis, inositol phosphate metabolism, and O-glycan metabolism were upregulated in ASD (Table S14).

To further explore metabolic differences across other ASD conditions, such as Asperger’s disorder (AD) and Pervasive Developmental Disorder-Not Otherwise Specified (PDD-NOS), we identified the significantly altered metabolic pathways for both AD and PDD-NOS. A PCA biplot, generated based on -log10 adjusted p-values from differential pathway analysis, revealed distinct metabolic activities across the three disorders, ASD, PDD-NOS, and AD. In PDD-NOS, inositol phosphate metabolism was upregulated, while in AD, glycosphingolipid biosynthesis (lacto and neo-lacto series) was upregulated. Cholesterol biosynthesis pathways were downregulated in PDD-NOS, while nucleotide metabolism and phosphatidylinositol phosphate metabolism were downregulated in AD (Figure S4). In ASD, pathways such as the pentose phosphate pathway (PPP), pyruvate metabolism, mitochondrial fatty acid β-oxidation, amino acid metabolism, heme synthesis, and porphyrin metabolism were downregulated, while O-glycan metabolism and fatty acid activation were upregulated (Figure S4).

We also performed reporter metabolite analysis to identify dysregulated metabolites and their associated pathways. In line with our findings in the brain cortex, metabolites related to oxidative phosphorylation were significantly enriched among downregulated genes (Figure 7). Additionally, metabolites from mitochondrial β-oxidation, BCAA metabolism, glycine, serine, and threonine metabolism, and sulfur amino acid metabolism were identified as reporter metabolites (Table S15). These findings suggest a consistent pattern of hypometabolism across both brain and blood in ASD.

**Figure 7.**
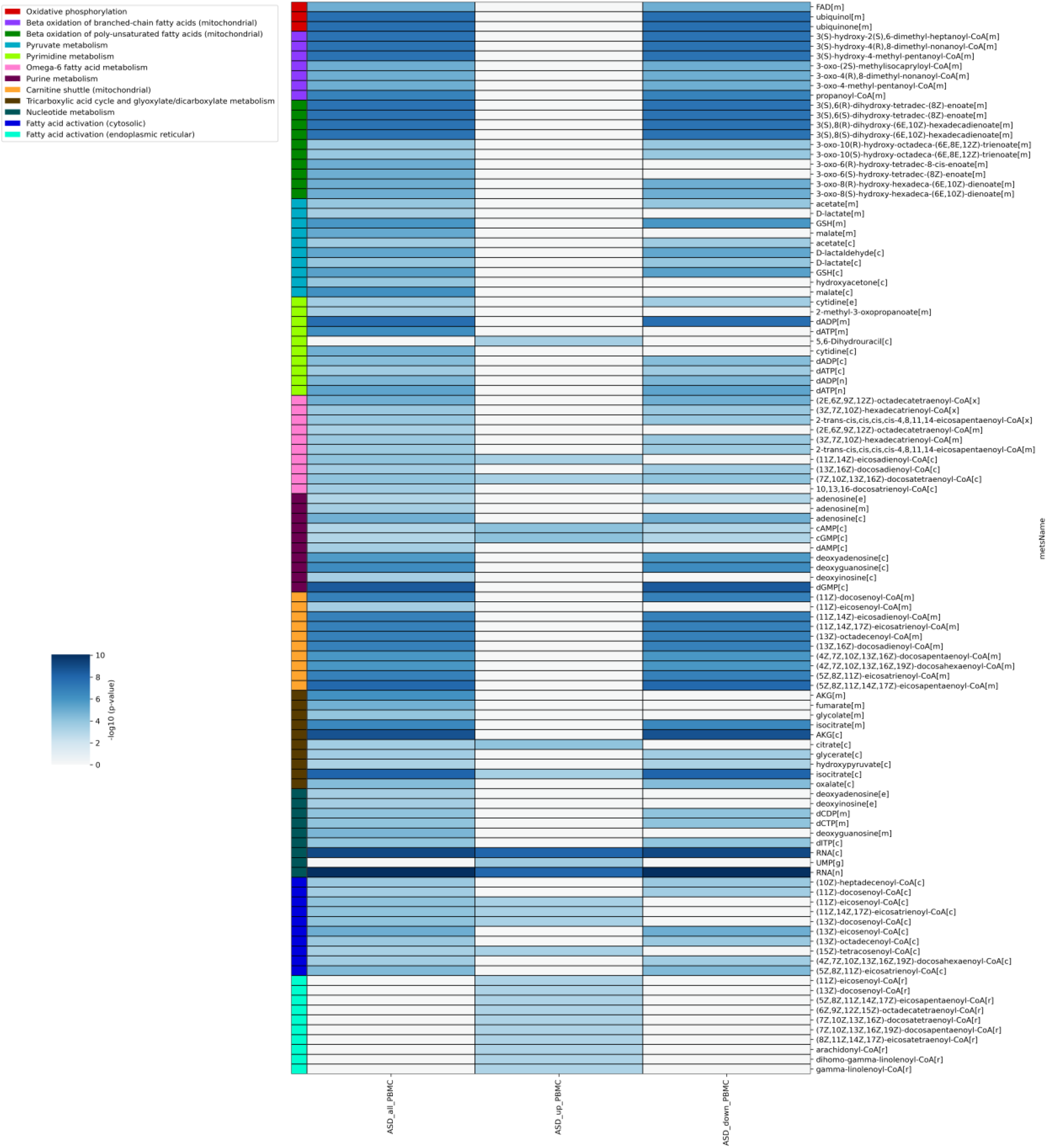
Heatmap of identified reporter metabolites that are altered in ASD compared to control in PBMC samples.

### Metabolic genes as diagnostic biomarkers

We investigated the diagnostic potential of metabolic pathway genes in identifying ASD using PBMC data. We computed the pathway activity score for metabolic pathways from the Human-GEM network for each sample. ROC analysis based on pathway scores revealed that pathways such as alanine, aspartate, and glutamate metabolism, inositol phosphate metabolism, and mitochondrial beta-oxidation of unsaturated fatty acids exhibited an AUC greater than 0.8 in the PBMC data (Table S16). Further, the differentially expressed metabolic genes between control and ASD were also used to perform the ROC analysis. A total of 78 metabolic genes showed an AUC greater than 0.8 (Table S17). These results were validated using both PBMC and leukocyte microarray data. In the PBMC dataset, 13 metabolic genes showed an AUC greater than 0.8. In the leukocyte transcriptome dataset, NDUFC1, CREBBP, DCXR, and ZDHHC17 showed an AUC greater than 0.7. Notably, CREBBP and ZDHHC17 displayed elevated expression in ASD compared to control and other spectrum disorders (Figure 8). CREBBP, which encodes CREB binding protein, is listed as an autism susceptibility gene in the SFARI database (https://gene.sfari.org/). ZDHHC17, involved in leukotriene metabolism, is essential for axon outgrowth and mediates the interaction between TrkA and tubulin during neuronal development [20].

**Figure 8.**
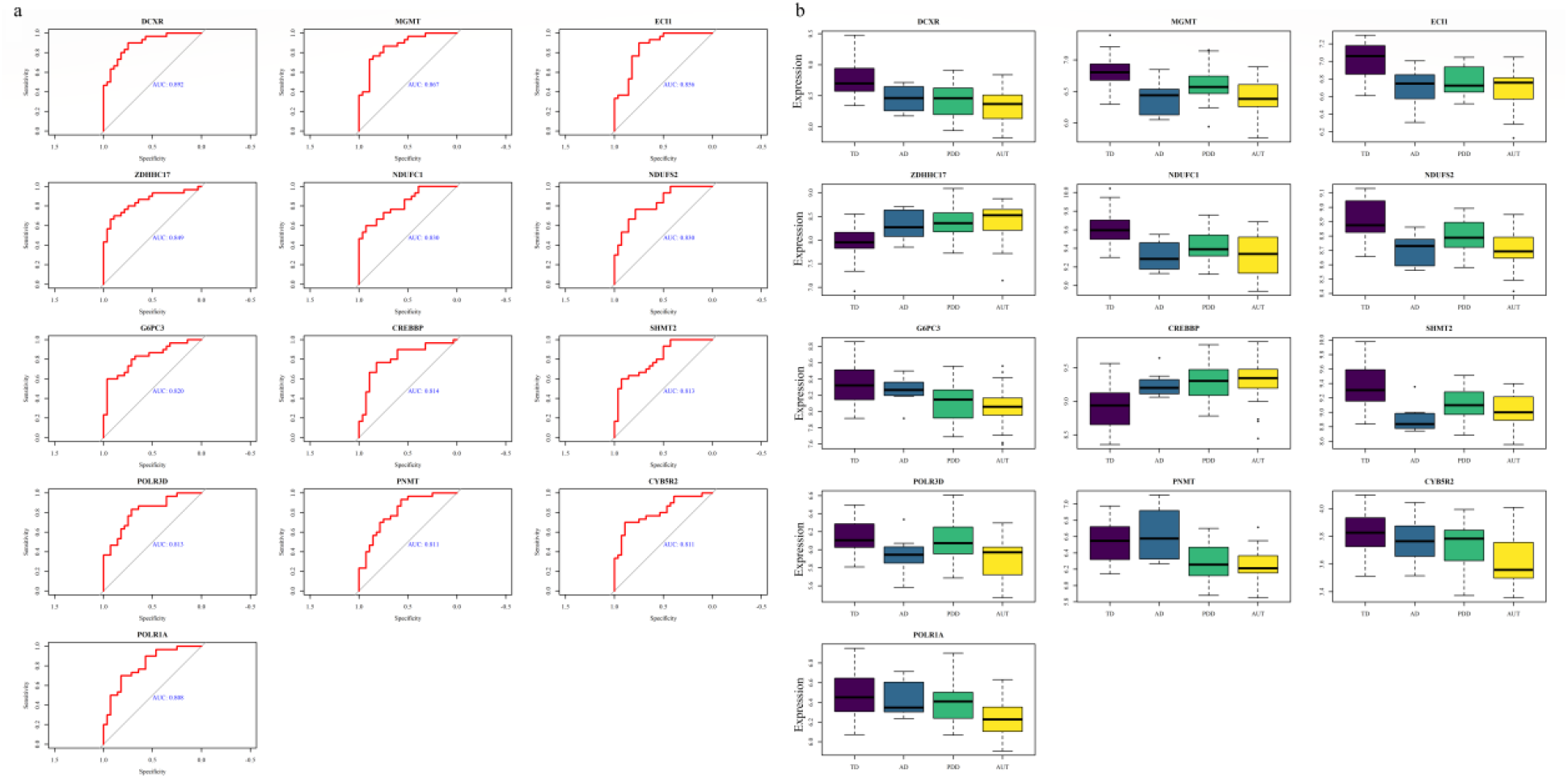
ROC curves of 13 differentially expressed genes with AUC > 0.8 in both PBMC transcriptome data (a) and expression levels of these 13 genes across ASD conditions.

Additionally, 15 differential expressed metabolic genes from PBMC data showed an AUC greater than 0.7 in the ASD cortex data (Figure S5, S6). ELOVL5, involved in the elongation of long-chain fatty acids, particularly those in the omega-6 and omega-3 families, exhibited increased expression in ASD compared to the control. SHMT2, a gene encoding serine hydroxymethyltransferase, also showed an AUC greater than 0.7. SHMT2 plays a crucial role in one-carbon metabolism, converting serine and tetrahydrofolate (THF) into glycine and 5,10 methylene THF. Impaired SHMT2 expression may lead to mitochondrial dysfunction, reduced cellular proliferation, and energy deficits, potentially affecting neural development and synaptic function.

## Discussion

We investigated the metabolic pathway activity in ASD using transcriptome data obtained from the brain cortex and blood samples. We identified key metabolic pathways, genes, and metabolites that may serve as potential biomarkers for ASD. Our findings highlight that critical metabolic pathways were downregulated in ASD. We observed the downregulation of energy metabolism pathways, including glycolysis/gluconeogenesis, pyruvate metabolism, the TCA cycle, and oxidative phosphorylation in both ASD brain cortex tissue and single-nucleus. This reduction in glucose metabolism suggests hypometabolism, a common feature in various neurological disorders, including Alzheimer’s and Parkinson’s diseases [21]. PET studies in ASD patients have shown hypometabolism in areas such as the parietal lobe, frontal premotor and eye-field areas, and the amygdala [22].

Glycolysis is crucial for membrane-associated processes like ion transport and is essential for maintaining neuronal inhibition through the phosphorylation of GABAA receptors, which regulate neurotransmission and synaptic activity [21]. The downregulation of the TCA cycle corroborated with alteration in metabolites such as citrate, malate, and oxaloacetate in the ASD brain. Disruptions in the TCA cycle can result in adverse neurodevelopment in affected individuals (Orozco et al., 2019). Alanine, aspartate, and glutamate metabolism, one step away from the TCA cycle, were also downregulated in the ASD brain. This disruption can impair the malate-aspartate shuttle (MAS), which is essential for transporting reducing equivalents (NADH) into the mitochondria [23]. Dysfunctional MAS can lead to ATP shortages in neurons, oxidative stress, and glutamate imbalances, all of which could contribute to neurodevelopmental deficits [24, 25]. Downregulation of oxidative phosphorylation in ASD may impair development since the developing brain has a high energy demand for processes like neuronal proliferation, migration, and synaptogenesis (formation of synapses) (Rae et al., 2024). We found that the ROS detoxification pathway and glutathione metabolism were dysregulated in the ASD brain, pointing to oxidative stress associated with ASD. This dysregulation may impair the ability to neutralize ROS, thereby exacerbating neuronal damage and contributing to ASD symptoms [26].

Our analysis also revealed increased mitochondrial β-oxidation in the cortex, indicative of altered fatty acid metabolism. While fatty acid oxidation can increase following brain injury, its upregulation in ASD may disrupt the balance between fatty acid synthesis and oxidation required for neurogenesis, potentially affecting neurodevelopmental outcomes [23]. Metabolites from omega fatty acid metabolism were identified as reporter metabolites, which are crucial for brain development and function. Abnormalities in omega fatty acid metabolism could impair the growth and functioning of neurons and their connections, contributing to neurodevelopmental issues in individuals with ASD [27].

Cholesterol metabolic pathways were also disrupted in ASD. Cholesterol plays a vital role in neurosteroid synthesis and synaptic formation. Notably, individuals with ASD exhibited significantly lower cholesterol levels, which are essential for maintaining neuronal health [28]. Altered cholesterol and bile acid metabolism have been implicated in Alzheimer’s disease [29]. Prostaglandin biosynthesis was upregulated in the ASD brain. Prostaglandin levels can influence neuronal migration and connectivity [30]. While certain prostaglandins provide neuroprotection, others can overstimulate neurons, potentially contributing to ASD symptoms such as hyperactivity and sensory sensitivities [31]. Elevated levels of PGE2 in the plasma of ASD patients have been linked to glutamate excitotoxicity and reduced GABA levels, potentially affecting social behavior [32]. GPI anchor synthesis, which aids in anchoring various proteins to the cell membrane, showed marked downregulation. Mutations in genes involved in GPI-anchor biosynthesis can lead to rare genetic disorders characterized by intellectual disability, seizures, and other neurological symptoms [33]. Deficiencies in GPI-anchored proteins may disrupt neuronal migration, synaptogenesis, and other developmental processes critical to ASD [34].

Reporter metabolite analysis identified metabolites in one-carbon metabolism in the ASD brain, including tetrahydrofolate (THF), 5,10-methenyl-THF, and 5,10-methylene-THF, asreporter metabolites. One-carbon metabolism supports DNA synthesis, repair, and methylation, which is essential for neuronal development [35]. THF facilitates the conversion of homocysteine to methionine, which is crucial for normal methylation patterns, while 5,10-Methenyl-THF and 5,10-Methylene-THF contribute to purine synthesis for DNA replication and repair. Disruptions in this pathway may lead to altered gene expression and impaired cellular function, which are often observed in neurological disorders [35]. Notably, the metabolic enzyme methyltetrahydrofolate dehydrogenase was differentially expressed in the ASD brain.

Vitamins B9 (folic acid), B1 (thiamine), B6, and B2 (riboflavin) were altered in ASD at the pathway and metabolite levels. Vitamin B6 is important for synthesizing neurotransmitters, including GABA, serotonin, dopamine, and noradrenaline, all showing impairment in ASD [36]. In rats, Vitamin B6 deficiency induces autism-like behaviors, marked by social deficits, repetitive behaviors, and altered GABAergic signaling and autophagy in the hippocampus [37]. Thiamine deficiency in ASD can impair brain function, contributing to symptoms such as impaired cognitive function and behavioral issues [38]. In ASD, riboflavin may play a protective role by reducing oxidative stress and supporting mitochondrial health [38]. It supports neurodevelopment processes, such as myelin synthesis.

Porphyrin metabolism, heme synthesis, and degradation were dysregulated in ASD, indicating potential disturbances in iron metabolism. We identified biliverdin, bilirubin, Fe2+, heme, carbon monoxide (CO) as reporter metabolites. Impaired heme synthesis can lead to oxidative stress, mitochondrial dysfunction, and disrupted neurotransmitter production, all of which are essential for proper neurological function. Iron, a key element in myelination and neurotransmitter production, is essential for brain development. Imbalances in iron levels can lead to significant developmental disruptions. Iron deficiency is reported in children with ASD [39]. Elevated levels of specific porphyrins have been observed in the urine samples of ASD (Kern et al., 2014). Altered porphyrin metabolism in ASD may also be linked to vitamin B6 deficiency, hyperoxalemia, hyperhomocysteinemia, and hypomagnesemia [40]. We observed that HMOX1, a gene encoding heme oxygenase-1 (HO-1), was upregulated in the ASD brain.

HO-1 is essential for maintaining redox balance, facilitating proper neuronal and glial maturation, and modulating synaptic activity. HO-1 protects against oxidative stress by degrading heme into biliverdin, iron, and CO. However, overexpression of HO-1 might lead to mitochondrial dysfunction, iron sequestration, and increased oxidative stress [41].

We observed a consistent pattern of hypometabolism in the PBMC of ASD patients, mirroring findings in the brain. Energy metabolism pathways, including glycolysis, the TCA cycle, and oxidative phosphorylation, showed reduced activity in ASD compared to control. Mitochondrial β-oxidation of fatty acids and the carnitine shuttle were also downregulated in ASD. In contrast, inositol phosphate metabolism was upregulated in ASD, helping to distinguish it from the control group with an AUC of 0.83, suggesting its potential as a diagnostic marker. Liu et al., (2024) also reported associations between inositol phosphate metabolism and ASD in their metabolomics study of autistic children [42]. Chondroitin and heparan sulfate biosynthesis was also upregulated with an AUC of 0.72. These glycosaminoglycans may influence neuronal migration, polarization, and the balance between neurogenesis and gliogenesis [43]. We identified metabolic genes that could serve as potential biomarkers for ASD, validating these findings across PBMCs and leukocytes.

Our work provides a comprehensive knowledge of metabolic pathway activity and potential biomarkers in individuals with ASD. The observed pattern of hypometabolism in both the brain and blood highlights its link to neurodevelopmental delays in ASD. Cross-disorder analysis revealed that these metabolic changes are more pronounced in ASD compared to SCZ, BD, and MDD. Further validation with larger cohorts is needed to explore potential therapeutic strategies.

## Author Contributions

Conceptualization: PKV; Methodology: RB; Formal analysis and investigation: RB, DS, AA; Writing - original draft preparation: RB; Writing - review and editing: RB, DS, AA, PKV; Funding acquisition: P KV; Supervision: PKV.

## Acknowledgments

This work was supported by iHUB-Data, International Institute of Information Technology, Hyderabad, India. The funding body has no role in the study design and analysis.

